# Bx42 directs neural stem cell exit from quiescence through Prospero

**DOI:** 10.64898/2026.08.11.744202

**Authors:** Nicole Losurdo, Uchechukwu E. Mgbike, Miranda Dietze, Isabella J. Hixson, Adriana Bibo, Xiao Mao, Nichole Link

## Abstract

Microcephaly is a rare neurodevelopmental disorder characterized by a severely reduced head and brain size in children and is accompanied by a myriad of debilitating side effects including cognitive and developmental impairments. While many cases of microcephaly arise from genetic mutations, the molecular mechanism linking variants to disease remain poorly understood. We previously identified *Bx42* as a microcephaly-causing gene from a patient-informed study using human brain organoid modeling and functional studies in *Drosophila melanogaster*. Here, we demonstrate that loss of *Bx42* leads to microcephaly by reducing neural stem cell proliferation early in development. Mechanistically, our data suggest that Bx42 promotes the transition from quiescence to proliferation through regulation of the neural differentiation factor Prospero. Reduction of Bx42 results in reduced neural stem cell division due to a prolonged quiescent state. Importantly, we demonstrate that two patient-derived variants in the human ortholog, SNW1, are nonfunctional or hypomorphic, providing strong evidence that these variants are pathogenic and causative of microcephaly. Together, our findings define a previously unrecognized role for the Bx42/SNW1 pathway in regulating neural stem cell activation and brain growth, offering new mechanistic insight into the pathogenesis of genetically driven microcephaly.

## Introduction

Microcephaly is a severe neurodevelopmental disorder that affects 8.7 out of 10,000 births in the United States and is characterized by a child’s orbitofrontal circumference measuring two or more standard deviations below the mean compared to individuals of the same age and sex (1–4). This rare disorder is often accompanied by comorbidities such as intellectual disability, seizures, motor and sensory impairments, and a reduced lifespan. Due to the severity of these neurological conditions, patients with microcephaly often need lifelong care. Many microcephaly cases have no known cause (3), and identifying which genetic mutations lead to microcephaly enhances our knowledge of understudied neurodevelopmental pathways, improves diagnostics for neurodevelopmental diseases, and promotes the possibility of therapeutic development.

Many characterized microcephaly-causing variants are in genes that regulate cell cycle progression and mitotic spindle orientation (4). Proper control of neural progenitor proliferation is essential for balanced brain growth (5), so neural stem cells (NSC) are one of the major cell types often affected in microcephalic brains (1). Much of the early work investigating NSC function used the *Drosophila* neuroblast as a model system (6–15). Due to these rigorous investigations, neuroblast division patterns and temporal gene regulation are well-known and characterized, making it an excellent model for studying novel gene functions. Further, human disease genes are often conserved between humans and flies, and functional analyses of patient variants are often efficacious in the fly (16–18). Together, this positions the *Drosophila* neuroblast as a key NSC model to study the function of novel human microcephaly genes.

NSCs can self-renew and produce differentiating daughter cells, and this balance is influenced by cell-fate determinants including transcription factors and signaling molecules (19). Through carefully controlled transcriptional regulation and localization of downstream targets, cell fate is balanced to promote either division or neuronal cell-fate specification (20). Changes in expression levels or asymmetric subcellular localization of important cell-fate determinants can drive the NSC either towards self-renewal, differentiation, or quiescence, a suspended state with no differentiation nor proliferation (21,22). While self-renewal is essential to maintain the NSC population and differentiation produces mature cell types such as neurons or glia, the quiescent state protects the cell from damage (23) and allows for neurogenesis at later life stages (24). When cell-fate determinants are disrupted, the NSC population can be lost via premature differentiation or a failure to exit quiescence, driving insufficient neuronal production and an overall reduction in brain size. These determinants could be defined as stemness factors, and we hypothesize microcephaly variants could affect these important genes.

We previously identified the fly gene *Bx42* from a microcephaly patient-informed investigation. *Bx42* and its human ortholog, *SNW1*, are components of the spliceosome and function as transcriptional regulators in some contexts. *Bx42* is necessary for early embryogenesis in flies since ubiquitous *in vivo* RNAi knockdown in the embryo results in dorsal closure failure, a hypotrophic nervous system, and lethality at embryonic stage 14 (25,26). Our previous work showed that *Bx42* reduction specifically in NSCs leads to a significant reduction in brain size and an absence of proliferation at the third-instar larval stage (27). We also showed that heterozygous mutations in human *SNW1* resulted in reduced organoid growth, indicating *Bx42/SNW1* is necessary for brain development. However, the functional consequences of SNW1 variants found in microcephaly patients have not been tested *in vivo*. Furthermore, the cellular mechanisms of Bx42 function in NSCs and the downstream pathways that influence NSC proliferation are unknown.

In this study, we find that proliferation defects recorded with reduction of *Bx42* are initiated shortly after larval hatching. At this time, NSCs normally enter a programmed quiescent phase but after 24 hours, promptly move into an active proliferation period through the remainder of larval development (22). We find that NSCs with reduced Bx42 cannot exit quiescence efficiently, leading to a loss of stem cells, decreased stem cell proliferation, and reduced brain size. The stem cell’s ability to enter and exit quiescence is regulated by the presence or absence of nuclear localization of the cell fate determinant Prospero (Pros) (28). In larvae with reduced Bx42, we find a significant increase in NSCs with nuclear Pros through all developmental stages.

Reducing Pros in NSCs can rescue phenotypes due to loss of Bx42 including brain size and stem cell phenotypes, showing that disrupted Pros drives NSC phenotypes with loss of Bx42. Together, our data show that Bx42 regulates NSC division, and thus brain size, through regulation of Pros expression, and Bx42 is necessary to maintain the propensity for a NSC to divide. Finally, we functionally test two microcephaly patient mutations *in vivo.* While wild type human SNW1 can rescue defects due to loss of Bx42, mutations found in microcephaly patients act as hypomorphs or loss-of-function, failing to rescue microcephaly phenotypes caused by Bx42 knockdown. Together, our data show that Bx42 is a critical factor that regulates the ability of NSCs to divide via Pros and establishes SNW1 variants as pathogenic.

## Results

### Bx42 is necessary for brain growth and NSC proliferation throughout larval development

We previously showed that knockdown of *Bx42* in NSCs leads to reduced brain lobe volume, fewer central brain NSCs, and a complete loss of proliferation at the late third-instar stage (27). To understand why loss of *Bx42* causes these phenotypes, we characterized stem cell behavior through key development timepoints with *in vivo* RNAi using the GAL4-UAS system and *inscuteable-GAL4* (*insc-GAL4*) for stem cell expression. NSCs are established during embryogenesis where they proliferate to produce the embryonic nervous system (29). At the end of embryogenesis, stem cells enter a state of quiescence (30) lasting ∼24 hours after larval hatching (ALH)(31) when animals begin eating. From this point in development, NSCs exit quiescence and proliferate continuously until pupariation (32). We assessed brain volume and NSC proliferation at 24 hours after larval hatching (L1 stage) using immunostaining for the NSC marker Deadpan (Dpn, magenta) and proliferation marker phospho-Histone H3 (pHH3, white). *Bx42* knockdown in NSCs caused a significantly reduced brain lobe volume (Figure 1A-C) and a reduction in dividing Dpn+ cells marked by pHH3 (Figure 1D-E, G), mirroring phenotypes found later in development (L3 stage (27)). However, the total number of Dpn positive cells did not significantly differ between *Bx42* knockdown and control animals (Figure 1F), suggesting Bx42 does not influence stem cell number at this early timepoint. Our data show that brain volume and proliferation defects start early.

**Figure 1.**
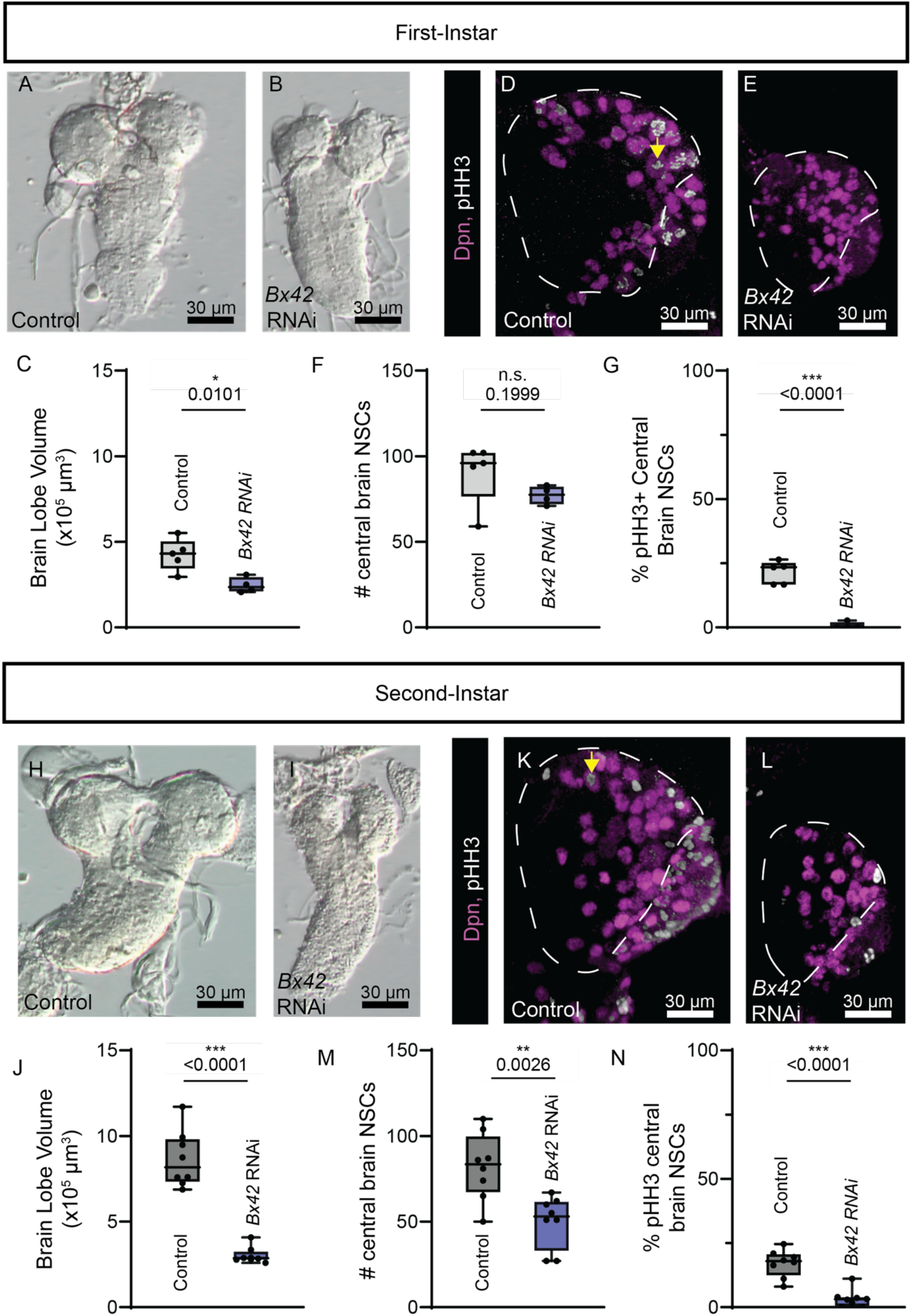
Reduction of *Bx42* in NSCs significantly reduces brain lobe volume and NSC proliferation. (A-B) Contrast images of first-instar larval brains taken 24 hours after larval hatching from knockdown in NSCs (*insc-GAL4*). (C) *Bx42* knockdown in NSCs results in significantly smaller brain lobe volume compared to *EGFP* RNAi (control, independent t-test: *t* = 3.495, *df* = 7, *p* = 0.0101, *n* = 4-5). (D-E) Confocal images of first-instar larval brain lobes stained with Deadpan (magenta, NSCs) and pHH3 (white, mitotic cells). Yellow arrow indicates a dividing NSC. (F) *Bx42* knockdown in NSCs does not significantly differ in the number of central brain NSCs compared to control at the first-instar larval stage (independent t-test: *t* = 1.415, *df* = 7, *p* = 0.1999, *n* = 4-5). (G) *Bx42* knockdown in NSCs resulted in a complete loss of mitotic central brain NSCs compared to the control at the first-instar larval stage (independent t-test: *t* = 8.849, *df* = 7, *p* < 0.0001, *n* = 4-5). (H-I) Contrast images of second-instar larval brains from *Bx42* knockdown with *insc-GAL4*. (J) *Bx42* knockdown in NSCs results in significantly smaller brain lobe volume compared to *EGFP* RNAi (control) at the second-instar stage (independent t-test: *t* = 9.268, *df* = 14, *p* < 0.0001, *n* = 8). (K-L) Confocal images of second-instar larval brain lobes stained with Deadpan (magenta) and pHH3 (white). (M) *Bx42* knockdown in NSCs had significantly fewer central brain NSCs than the control at the second-instar larval stage (independent t-test: *t* = 3.664, *df* = 14, *p* = 0.0026, *n* = 8). (N) *Bx42* knockdown in NSCs results in significantly less mitotic central brain NSCs compared to the control at the second-instar larval stage (independent t-test: *t* = 6.098, *df* = 14, *p* < 0.0001, *n* = 8).

To determine whether proliferation defects are maintained into later stages with loss of Bx42, we assessed stem cell numbers and proliferation in the second-instar stage (L2). Brains from animals with reduced *Bx42* were significantly smaller compared to controls (Figure 1H-J) like observations at both earlier (L1) and later (L3) developmental time points. *Bx42* knockdown animals also had significantly fewer Dpn positive central brain NSCs (Figure 1K-M), establishing that stem cell loss arises by the second instar stage. While there was a significant reduction in the percentage of proliferating central brain NSCs (pHH3 +, Dpn +, Figure 1N), some cells did enter mitosis. For a brief developmental window during L2 to early L3, some Dpn positive cells lacking Bx42 can proliferate, but their low abundance cannot compensate for early defects affecting stem cell division.

### Bx42 prevents NSC nuclear Prospero expression

We hypothesized that many stem cells are failing to exit the quiescent state due to the absence of proliferation at 24 hours after larval hatching and low proliferation rates at the L2 stage. The NSC’s ability to enter and exit quiescence is regulated by intrinsic cues including levels of nuclear Pros. A transient increase in the levels of Pros in the nucleus pushes the cell towards quiescence, while no nuclear Pros ensures NSCs can enter mitosis (8,10,13,28). To determine whether Bx42 affected Pros localization or expression levels, we checked for the presence of nuclear Pros in Dpn + cells across development using immunostaining. In control L1 animals, less than 5% of Dpn + cells contained nuclear Pros, and no nuclear Pros was detected in L2-L3 NSCs (L1, Figure 2A-A’’’; L2, Figure 2C-C’’’; and L3, Figure 2E-E’’’). These results agree with previous literature showing NSCs in these developmental stages are not normally in a state of quiescence (7,32).

**Figure 2.**
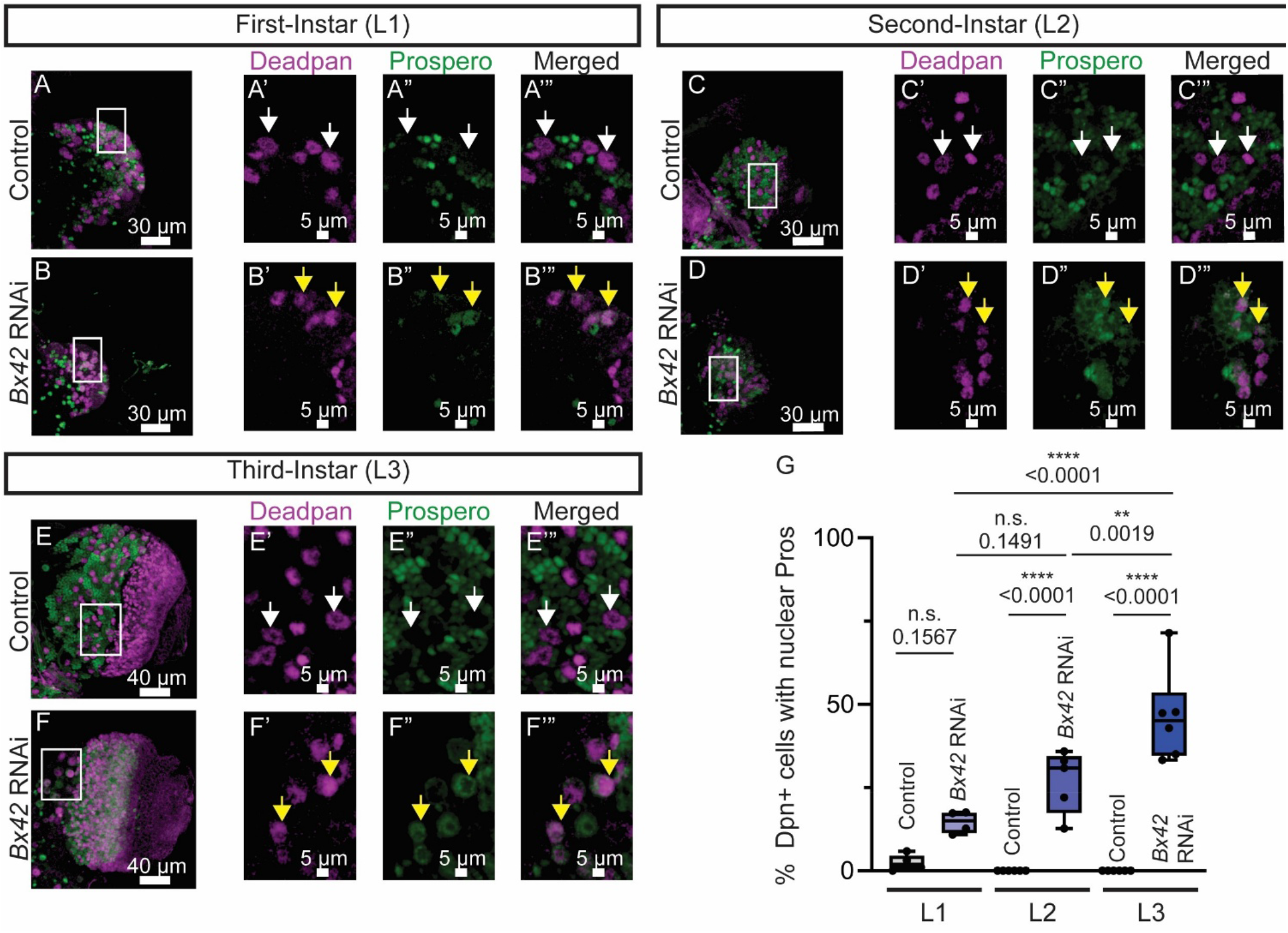
Bx42 prevents nuclear Pros in central brain NSCs throughout development. Confocal images of (A-B) first-instar 24 hours ALH, (C-D) second-instar, and (E-F) third-instar larval brain lobes. All brains were stained with Deadpan (Dpn, magenta, NSCs) and Prospero (Pros, green, early neurons), and knockdown was in NSCs (*insc-GAL4*). The white box indicates the zoomed-in region for panels. A white arrowhead indicates a NSC without nuclear Pros, and a yellow arrowhead indicates a NSC with nuclear Pros. (G) There was no significant difference in the percentage of nuclear Pros at the first-instar stage (One way ANOVA, p=0.1567). At both the second-(p<0.0001) and third-instars (p<0.0001), *Bx42* knockdown brains had significantly higher percentages of NSCs with nuclear Pros compared to controls. The percent of NSCs with nuclear Pros also significantly increased over time.

However, nuclear Pros was present in *Bx42* knockdown Dpn + cells in all three developmental stages (L1, Figure 2B-B’’’; L2, Figure 2D-D’’’; L3, Figure 2F-F’’’). By L2, the percentage of Dpn + cells with nuclear Pros was significantly greater than control (Figure 2G), and L3 animals contained a significantly higher percentage of nuclear Dpn +, Pros + cells than L2 animals, demonstrating that nuclear Pros increases over developmental time. Our results suggest that Bx42 negatively regulates Pros expression in the nucleus to maintain the proliferative stem cell state.

### Bx42 inhibits NSC quiescence

Since nuclear Pros precedes full NSC quiescence and can decrease once a cell is in the quiescent state (33), we decided to use additional methods to quantify quiescence. One morphological characteristic of quiescent NSCs is a singular cytoplasmic protrusion that connects to the neuropil (34–36). These protrusions are identifiable as long extensions of the cell cortex and visualized when the cortex is labeled. We chose to use Miranda immunolabeling since it localizes to the cortex in an interphase cell and is specific to NSCs. We stained brains at 0-2 hours ALH (Fig 3A-B), 24 hours ALH (Fig 3C-D), and during wandering L3 (Fig 3E-F) stage with Miranda (magenta) and Pros (green). At 0-2 hours ALH, ∼30% of control NSCs had protrusions (Figure 3A-A’, G), and by 24 hours ALH, this number significantly reduced to only 5% (Fig 3C-C’, G), consistent with published literature showing almost all NSCs exit quiescence by 24 hours ALH (30). No protrusions were identified in control brains at the L3 stage (Figure 3 E-E’, G), matching the observation that most NSCs are cycling at this stage. However, when *Bx42* was knocked down, consistently 40% of NSCs had protrusions across each developmental stage tested (Figure 3). These results were also significantly different than the number of protrusions in control brains at every timepoint (Figure 3G). The persistence of protrusions past 24 hours ALH indicates a failure of NSCs to exit quiescence, leading to a reduced number of mitotic NSCs and fewer differentiating progeny.

**Figure 3.**
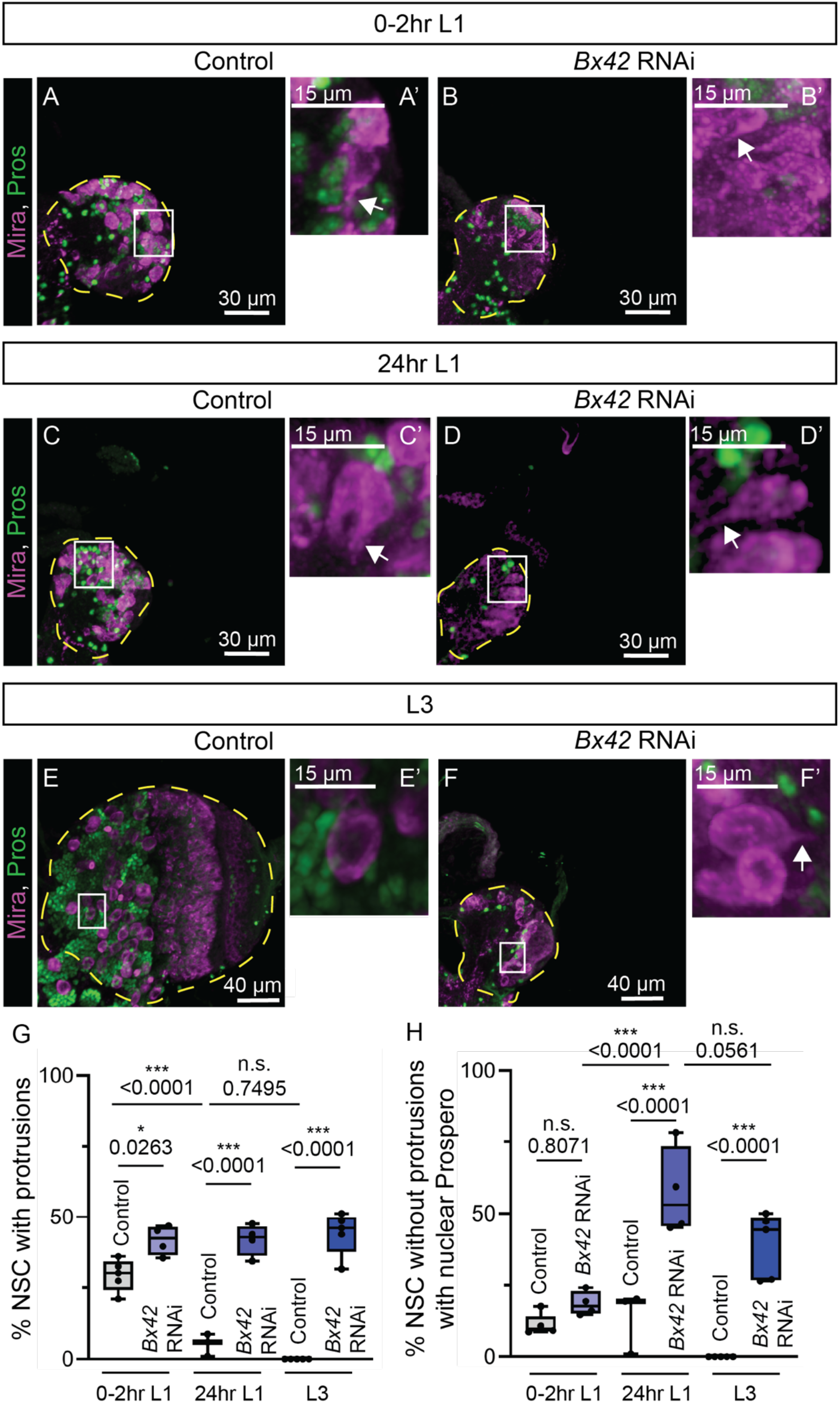
Loss of *Bx42* results in increased number of NSCs with a single protrusion throughout development. (A-F) Confocal images of *EGFP* RNAi (A, Control) or (B) *Bx42* RNAi 0-2 hours ALH first-instar (L1), (C-D) 24 hours ALH first-instar, and (E-F) third-instar (L3) larval brains stained with Miranda (magenta) and Pros (green). (A’-F’) Zoomed in images of white box regions showing individual NSCs. White arrow indicates a protrusion. (G) Percent of central brain NSCs with a singular protrusion. Control brains display a significant reduction of protrusions between 0-2 hours ALH and 24 hours ALH in the first-instar stages as expected for exit from quiescence (Tukey’s multiple comparisons ANOVA, p<0.0001). *Bx42* knockdown brains have a significantly higher percentage of protrusions at 0-2 hours ALH that maintains throughout development (Tukey’s multiple comparisons ANOVA, p = 0.0247 0-2hrs, p<0.0001 24hrs and L3). (H) Percent of stem cells without protrusions that have nuclear Pros. Control first-instar brains at both time points have low percentages of stem cells with nuclear Pros that is absent by the third-instar. *Bx42* knockdown brains have no differences in nuclear Pros in stem cells without protrusions at 0-2 hours ALH in the first-instar stage (Tukey’s multiple comparisons ANOVA, p = 0.7883). By 24 hours ALH and in the 3^rd^ instar stage, *Bx42* knockdown brains have significantly more stem cells without protrusions and with nuclear Pros (Tukey’s multiple comparisons ANOVA, p<0.0001).

We then looked at the combined phenotypes of protrusions and nuclear Pros to determine if they were marking the same cells. As expected, some NSCs with protrusions had nuclear Pros, indicating they are quiescent. We observed that almost 60% of the protrusion negative NSCs have nuclear Pros at 24 hours ALH (Fig 3H). While low levels of nuclear Pros activate quiescence in NSCs, high levels initiate genes necessary for terminal differentiation(28,33). Since Pros is necessary to transition a cell between quiescence, differentiation and self-renewal, and protrusions are markers of quiescent cells, we hypothesize that NSCs without protrusions but nuclear Pros + could be terminally differentiating in addition to entering quiescence. Conversely, around 40% of NSCs are already quiescent.

### Bx42 prevents nuclear Prospero in NSCs to maintain stemness

Bx42 likely regulates many targets by impacting splicing or transcription. Increased nuclear Prospero could be a direct effect of reducing Bx42, or a downstream consequence of cells undergoing quiescence driven by another Bx42 target. To differentiate between these two mechanisms, we performed a genetic epistasis test between *Bx42* and *pros*. To assess this, we knocked down both *pros* and *Bx42* in NSCs in the same animal and compared phenotypes to knockdown of *pros* or *Bx42* alone. Knockdown of *pros* alone had a significantly smaller brain size compared to control (Figure 4A-B, E). To control for possible GAL4 dilution effects, we introduced an irrelevant RNAi line targeted to *EGFP* together with *Bx42* knockdown. Knockdown of both *EGFP* and *Bx42* in the same animal with *insc-GAL4* resulted in significantly smaller brains (Figure 4C, E), phenocopying single knockdown of *Bx42* (27) and indicating no GAL4 dilution effect with two UAS lines.

**Figure 4.**
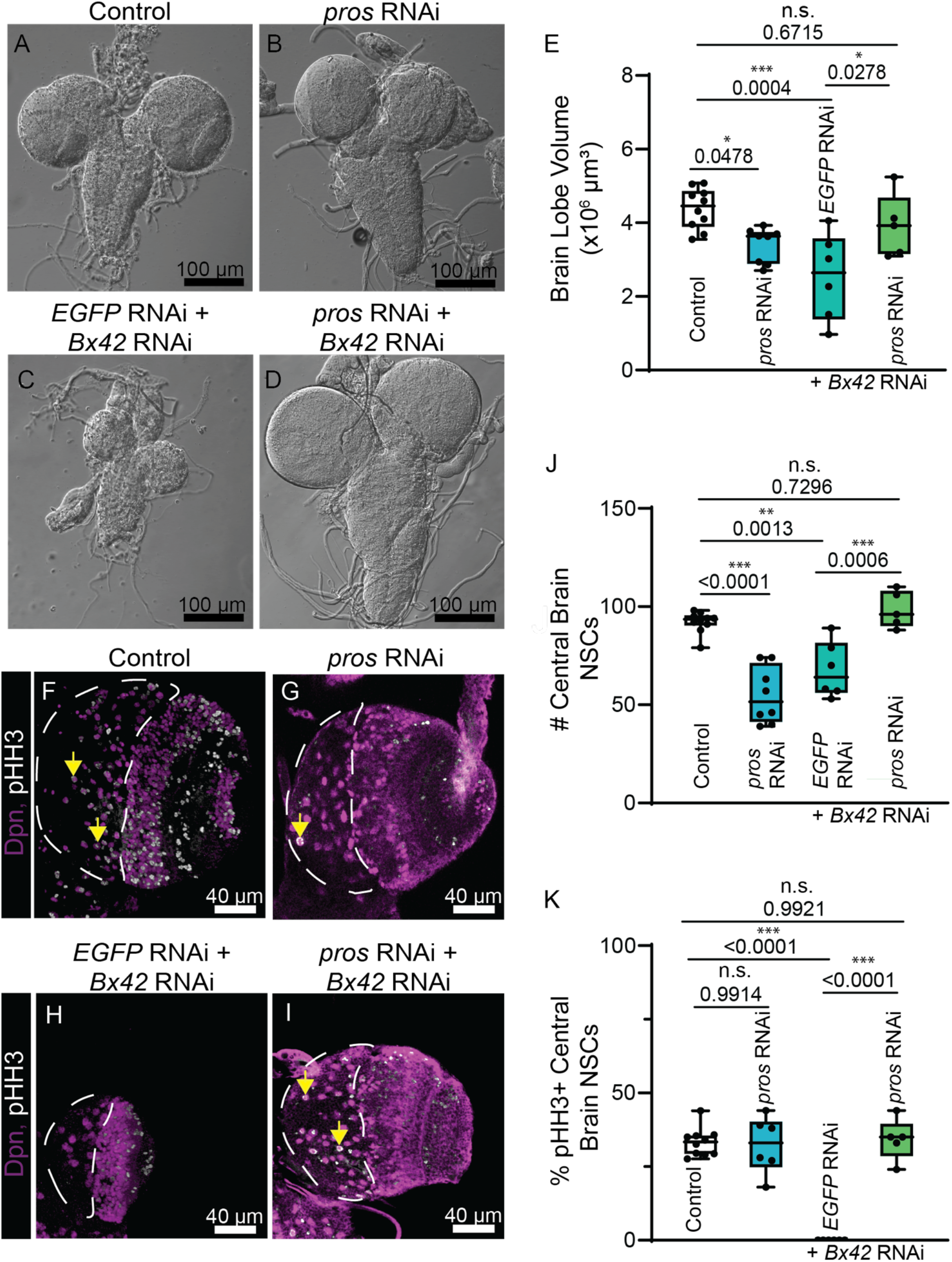
Reduction of *Pros* rescues *Bx42-*induced defects. Contrast images of third-instar larval brains from knockdown in NSCs (*insc-GAL4*). (A) Control (*EGFP* RNAi), (B) *pros* RNAi, (C) *EGFP* and *Bx42* RNAi, and (D) *pros* and *Bx42* RNAi. (E) Quantification of brain volume of brains from (A-D). Double knockdown of *pros* and *Bx42* is significantly larger than *EGFP* + *Bx42* double knockdown (Tukey’s multiple comparisons ANOVA: *p* = 0.0278) and did not differ from control (Multiple comparisons ANOVA: *p* = 0.6715), indicating full rescue of brain lobe volume. (F-I) Confocal images of late third-instar larval brain lobes stained with Deadpan (magenta, NSCs) and pHH3 (white, mitotic cells). The central brain region is outlined in white. A few pHH3+ cells are marked with yellow arrows. (F) Control (*EGFP* RNAi), (G) *pros* RNAi, (H) *EGFP* RNAi + *Bx42* RNAi, and (I) *pros* RNAi + *Bx42* RNAi. Quantification of (J) Central Brain NSC number and (K) dividing NSCs in brains represented in (F-I). Double knockdown of *EGFP* and *Bx42* resulted in (J) significantly fewer central brain NSCs than control and (K) no dividing NSCs, similar to *Bx42* knockdown alone (Tukey’s multiple comparisons ANOVA: *p* = 0.0013 and p<0.0001). Double knockdown of *pros* and *Bx42* rescues (J) central brain NSC number and (K) the percentage of dividing NSCs (Tukey’s multiple comparisons ANOVA: *p* = 0.0006 and p<0.0001), indicating full rescue.

However, double knockdown of both *pros* and *Bx42* resulted in brain volumes comparable to controls (Figure 4A, D-E) and significantly larger than Bx42 knockdown animals (Figure 4C, E) (27). These results show that Pros reduction can rescue Bx42-induced brain size defects and suggest that Prospero could be the driving factor of the stem cell deficits that lead to microcephaly with loss of Bx42.

We next assessed if reduction of Pros could also rescue NSC number and proliferation phenotypes due to loss of *Bx42*. *pros* knockdown alone reduces NSC number (Figure 4F-G, J) but did not affect the percentage of dividing NSCs compared to controls (Figure 4F-G, K). Double knockdown of *EGFP* and *Bx42* mirrors *Bx42* knockdown alone with a significant reduction in NSC number and a complete absence of mitotic NSCs (Figure 4H, J-K), again demonstrating GAL4 dilution effects are not a concern for our system. The double knockdown of *pros* and *Bx42* fully rescues both NSC number and percent NSC proliferation compared to Bx42 knockdown, and neither are statistically different than controls. Our results show that Pros reduction rescues *Bx42* knockdown stem cell phenotypes. Thus, Pros drives Bx42-induced phenotypes in the NSC during brain development, and it is likely that Bx42 directly suppresses Pros to maintain the NSC’s propensity to divide.

### Human SNW1 variants are hypomorphic or loss-of-function

We previously showed that human SNW1 and Bx42 are functionally conserved in brain development (27). Alignment of Bx42 and SNW1 proteins shows 60% conserved identity between amino acid residues.

Heterozygous deletions in *SNW1* result in reduced organoid growth (27), but patient variants were not functionally tested. In this study, we used *Drosophila* to test the function of microcephaly patient variants *in vivo*. Two variants from Ji et al., 2025 (27) were tested (Figure 5A): 1) c.182_187del, p.G61_G62del and 2) c.1235_1236insA, p.F412Lfs*17. All residues were similar or identical between human SNW1 and fly Bx42. Variants were generated in human UAS-human SNW1 constructs with a C-terminal HA tag.

**Figure 5.**
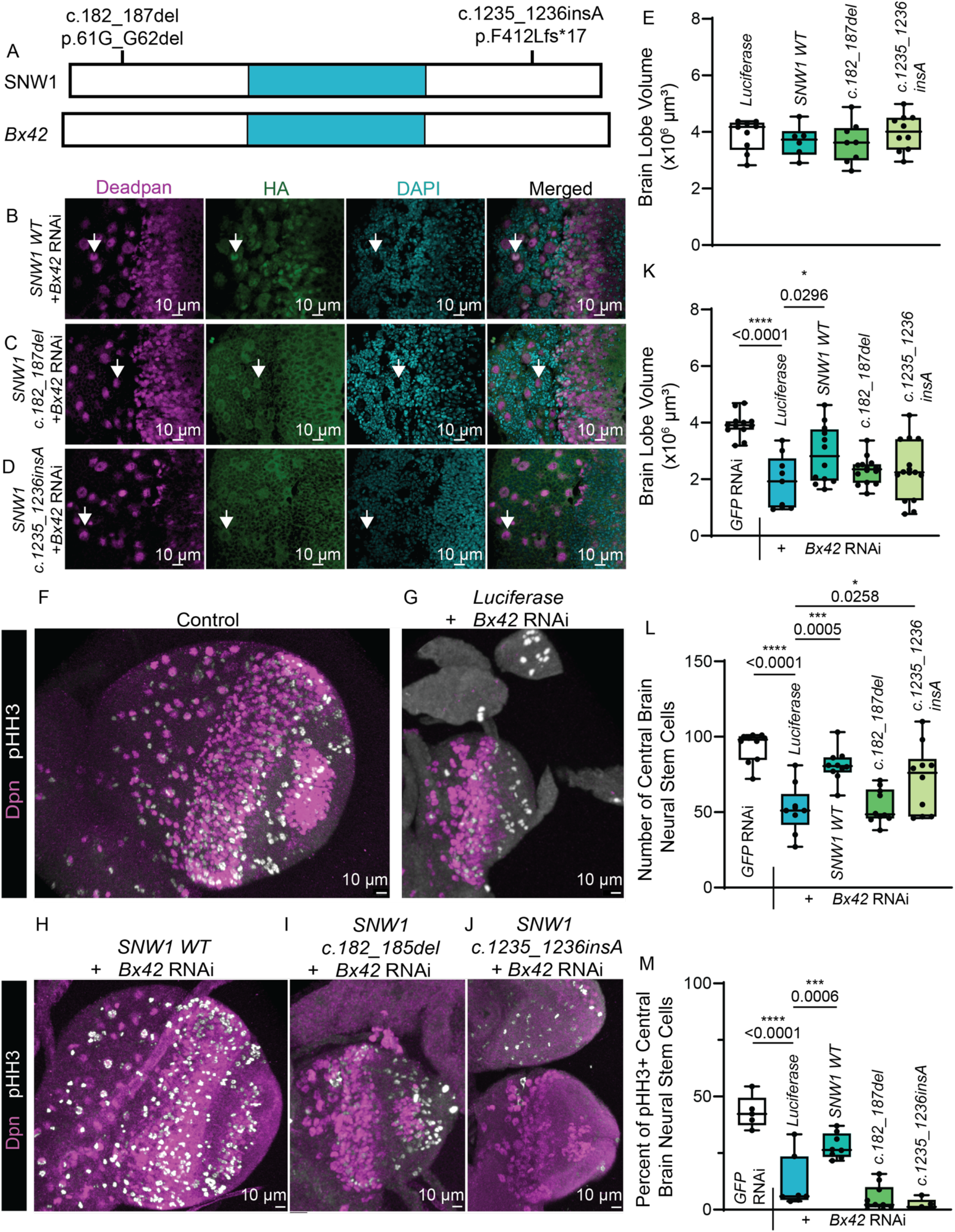
Human *SNW1* variants are hypomorphic or loss-of-function mutations. (A) Aligned Bx42 and SNW1 proteins with SNW1 variants tested. Blue indicates the SNW domain. (B-D) Fluorescent images of brains with expression of wild type or variant *SNW1* in NSCs (*insc-GAL4*). NSCs (Dpn, magenta), SNW1 (green), and DAPI in third-instar larval brains. Only *SNW1 WT* showed strong nuclear localization. (F) Expression of *SNW1* wild type and patient variants alone showed no difference in brain lobe volume compared to controls. (F-J) Fluorescent confocal images (partial z-stacks) of *SNW1* variants + *Bx42* RNAi in NSCs (*insc-GAL4)* stained for NSCs (Dpn, magenta) and proliferation (pHH3, white). (K) NSC expression of *Luciferase + Bx42* knockdown resulted in significantly smaller brain lobes than control (Sidak’s multiple comparisons ANOVA: *p* < 0.0001). Expression of wild type SNW1 rescued *Bx42* knockdown brain size but patient variants did not. (L) Number of central brain NSCs. *Luciferase + Bx42* knockdown resulted in significantly less NSCs than control (Sidak’s multiple comparisons ANOVA: *p* < 0.0001). Expression of wild type *SNW1* significantly rescued loss of *Bx42* (Sidak’s multiple comparisons ANOVA: *p* = 0.0011)*. SNW1 c.1235_1236insA* also rescued stem cell number but *c.182_187del* did not. (M) Percentage of proliferating NSCs in *Luciferase + Bx42* RNAi resulted in significantly less proliferating NSCs than control (Sidak’s multiple comparisons ANOVA: *p* < 0.0001). Expression of SNW1 wild type significantly rescued dividing NSCs but other variants did not (Sidak’s multiple comparisons ANOVA: *p* = 0.006).

We first verified expression stability of the wildtype and variant SNW1 constructs using HA immunostaining (Figure 5B-D). The SNW1 protein localizes to the nucleus as seen in the wildtype construct (Figure 5B). However, c.182_187del and c.1235_1236insA mutations show a dispersed HA expression that is both nuclear and cytoplasmic (Figure 5C-D), indicating that SNW1 localization is disrupted in these variants. We also tested whether expression of wild type or variant SNW1 in a wild type background caused gain-of-function or toxic overexpression effects using brain lobe volume as a readout. We found no differences in size when compared to a UAS-Luciferase control (Figure 5E). Over-expression of wild type and variant human SNW1 in *Drosophila* does not cause dominant or toxic effects.

We previously showed that the function of SNW1 is conserved with Bx42 using neurodevelopmental rescue experiments (27). Wild type human SNW1 could rescue brain volume defects, NSC number, and NSC proliferation rates when raised at 25°C (27). Here, we repeated the same experiment at higher temperatures (27°C and 29°C) to induce stronger phenotypes from *in vivo Bx42* RNAi. Very few animals emerged when crosses were reared at 29°C, so we assayed SNW1 variants at 27°C. At this temperature, loss of *Bx42* using RNAi again caused microcephaly that was rescued by expression of wild type human SNW1 (Figure 5F-K), and no GAL4 dilution effects were detected (*Bx42* RNAi + UAS-Luciferase). Since we validated conservation in multiple conditions, we co-expressed patient variants of SNW1 with *Bx42* RNAi and analyzed resulting brain size and NSC phenotypes. Neither variant was able to rescue small brain size when co-expressed with *Bx42* RNAi, suggesting that both variants are loss-of-function. Co-expression of UAS-*Luciferase* with *Bx42* RNAi also phenocopied the reduction of NSC number and proliferation (Figure 5N-O) seen in *Bx42* RNAi alone (27). Co-expression of wild type human SNW1 with *Bx42* RNAi significantly rescued NSC number (Figure 5H, L) and the percentage of dividing stem cells (Figure 5M), although not to wild type levels. Despite its inability to rescue brain lobe volume (Figure 5K), the *SNW1* mutant c.1235_1236insA partially rescued NSC number but not proliferation (Figure 5J, L, M). These data suggest that c.1235_1236insA is hypomorphic and retains some function. c.182_187del (Figure 5I) could not rescue brain size (Figure 5K), NSC number (Figure 5L), or NSC proliferation (Figure 5M), indicating it is likely a loss-of-function mutation. Together these data show that while *SNW1* mutants have varying levels of severity, all are pathogenic and able to cause microcephaly.

## Discussion

We investigated biological mechanisms linked to *Bx42/SNW1* associated microcephaly, or reduced brain size. Loss of Bx42 in flies and SNW1 in humans causes a small brain phenotype, and microcephaly patient variants in SNW1 act as loss-of-function or strong hypomorphs when tested using *Drosophila*. Loss of Bx42 impacts NSCs as early as 24 hours ALH with phenotypes that persist through larval development. NSCs remain in a non-dividing, quiescent state, driven by excessive Pros in the nucleus. Importantly, reduction of Pros suppresses brain volume defects caused by loss of Bx42, suggesting that Pros is downstream of Bx42 in a direct causal relationship. Our results show that Bx42 is a novel regulator of Pros and demonstrate that Bx42 is an important factor for maintaining the NSC’s proliferative capacity and other stemness qualities. As a result, this pathway could be linked to additional neurodevelopmental diseases that are the result of NSC defects in humans.

Bx42/SNW1 is a member of the spliceosome complex and has been shown to regulate splicing of multiple targets (27,37,38). In addition, the protein acts as a transcriptional co-activator or repressor, depending on what complex it binds (39). We predict that Bx42/SNW1 could have numerous downstream targets that affect development, and we initially anticipated a complex mechanism of action. However, our rescue experiments suggest that Pros is the main target of Bx42 in NSCs during brain development, showing that even splicing factors and transcriptional regulators can have very specific roles in distinct tissue types.

Since Pros is upregulated by loss of Bx42, we hypothesize that the normal function of Bx42 is to repress Pros transcription. It will be interesting to identify other Bx42 interaction partners during neurodevelopment to define proteins that cooperate with Bx42 to transcriptionally repress Pros.

Increased nuclear Pros could lead to two different scenarios: 1) premature differentiation and NSC loss or 2) a block from quiescence exit and no entry into the cell cycle. Either situation causes loss of dividing NSCs, a reduction of differentiating daughter cells, and eventual microcephaly. We document both loss of NSCs and progenitor cells that retain some NSC qualities, so we predict both scenarios occur in the developing brain. It is possible that elevated nuclear Pros has stage-specific effects, promoting differentiation in some cells and maintaining quiescence in others. The coexistence of both quiescent and differentiating NSC phenotypes suggest that precise regulation of Pros localization and protein level is required for the balance of stem cell maintenance, cell-cycle reentry, and differentiation during development. Lineage tracing tools could be useful to verify these hypotheses and define the contribution of these mechanisms to the observed microcephaly phenotypes with loss of Bx42.

The human ortholog of Pros is PROX1, which has a rich literature describing its importance during development (40–42). Interestingly, variants in PROX1 have not been definitively linked to neurodevelopmental disease. Hints of misexpression are associated with a KIF11-mediated form of microcephaly-lymphedema-chorioretinopathy syndrome (MLC), but in this scenario, PROX1 expression was decreased (43). Nevertheless, it will be important to investigate whether SNW1 directly impacts PROX1 expression in human models. If SNW1 loss increases nuclear PROX1 in human stem cells, PROX1 degradation could be tested for a therapeutic intervention in models of SNW1 neurodevelopmental disease. In addition, whether PROX1 is linked to additional neurological disorders is an open and intriguing question. Our findings raise the possibility that dysregulation of PROX1 activity, localization, or protein stability may represent an underappreciated mechanism contributing to defects in neural stem cell maintenance or brain growth. Studies examining PROX1 protein dynamics in patient derived models may provide additional insight into SNW1-associated microcephaly and other neurodevelopmental disorders.

Here, we identified a conserved pathway important for neurodevelopment and linked to human disease. By combining functional studies in *Drosophila* with human microcephaly patient data, we have identified a stemness factor important for maintaining stem cell proliferative potential. The presence of Bx42, and perhaps SNW1, is essential to preserve a self-renewing state that retains NSC numbers and produces differentiating progeny that populate the brain. Characterizing how SNW1 and its downstream effectors coordinate stem cell maintenance across species will improve our understanding of brain development but may also reveal new therapeutic opportunities for disorders associated with impaired neurogenesis or reduced brain growth. Our work using *Drosophila* demonstrates that the study of SNW1 function and characterization of its downstream targets have the potential to provide critical insights into stem cell biology and disease.

## ACKNOWLEDGEMENTS

We would like to thank members of the Link lab for suggestions and critical reading. We thank the Bloomington *Drosophila* Stock Center and Developmental Studies Hybridoma Bank for providing stocks and reagents. We acknowledge HSC Cell Imaging Core at the University of Utah for use of equipment. This work was supported by the National Science Foundation Graduate Research Fellowship Program (2139322) to NLosurdo.

## AUTHOR CONTRIBUTIONS

The number of experiments performed by each researcher was the method used for assigning the order of the 3 co–first authors. NLosurdo led experiments except for Prospero epistasis (UEM) and human rescue analysis (MD). IJH and AB assisted throughout. XM provided patient information. NLosurdo, UEM, MD, and NLink wrote the manuscript. NLink supervised the project design and revised the manuscript.

## Key Resources

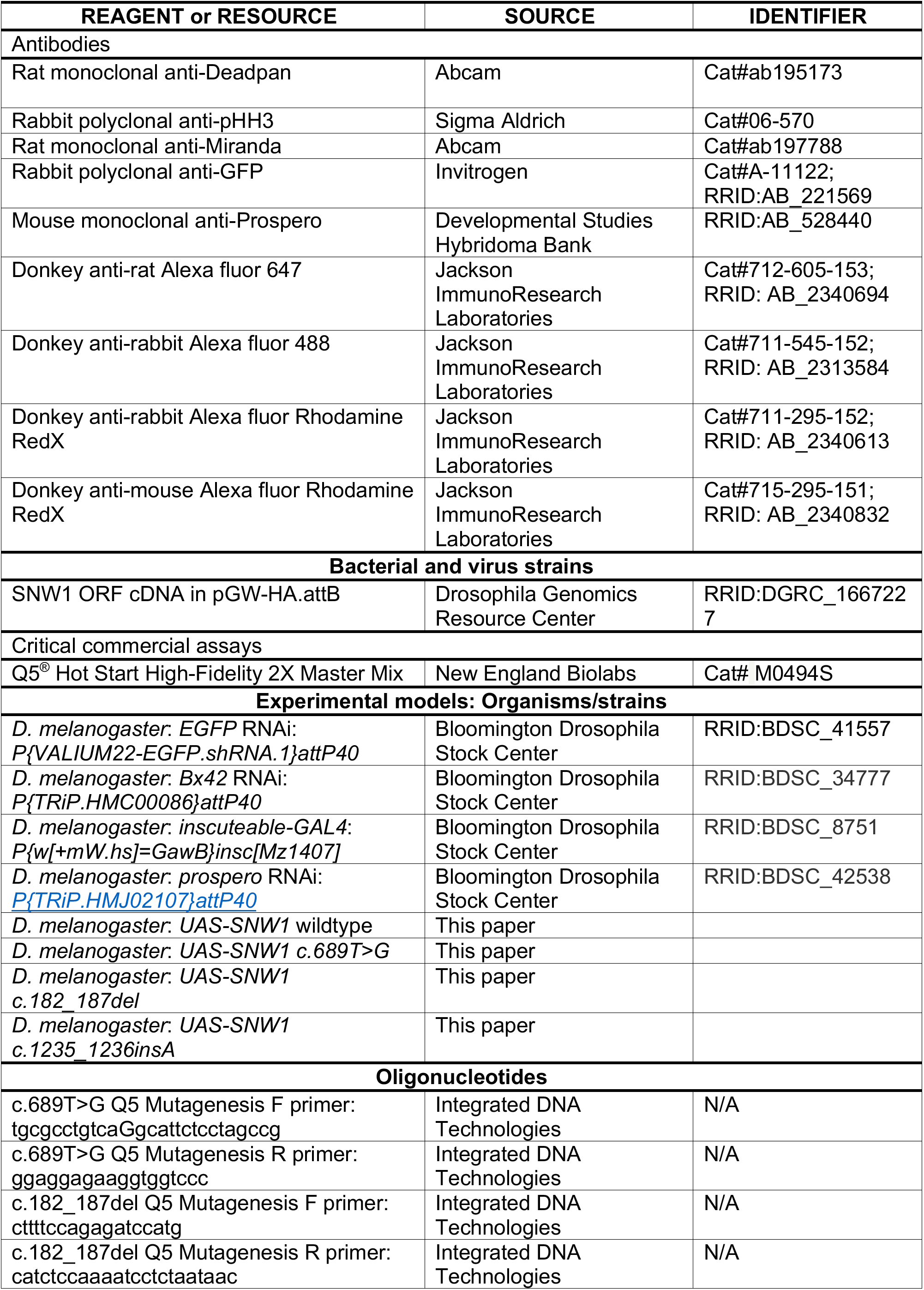
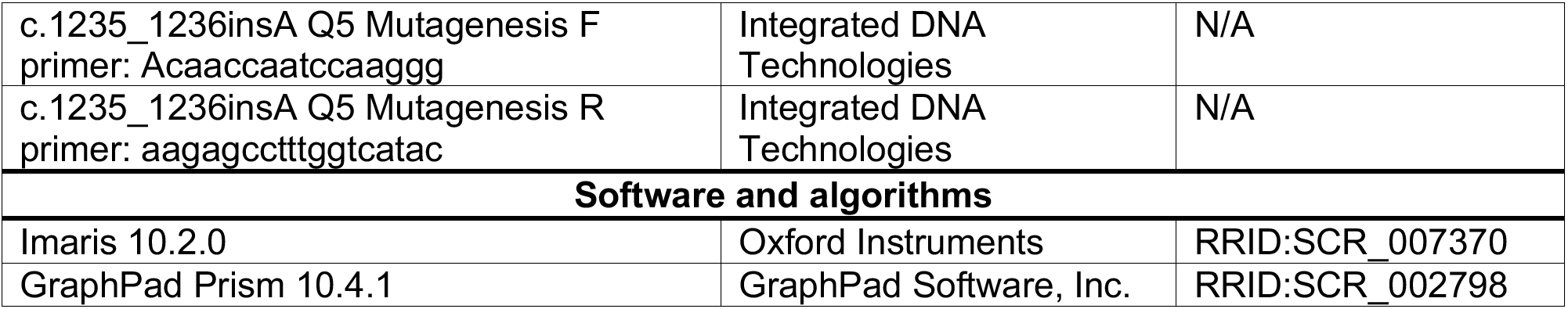

## EXPERIMENTAL MODEL AND STUDY PARTICIPANT DETAILS

### Fly lines and maintenance

The *Drosophila melanogaster* lines used in this study are listed in the key resources table. All stocks were raised at 25°C on Archon glucose formula food in wide vials. Most crosses were set at 29°C on blue food which was made by adding bromophenol blue to the glucose food. Rescue crosses were set at 27°C. Late third-instar larvae were selected for brain dissection based on extruding spiracles and gut clearance. The second-instar larval stage was identified by mouth hook morphology. The late first-instar larval stage was defined as 24 hours after larval hatching (ALH). Early first-instar larval stage was defined as 0-2 hours ALH.

## METHOD DETAILS

### Generating human variant flies

Wildtype human *SNW1* ORF cDNA hGUHO12880 (hGUHO12880 (DGRC Stock 1667227; https://dgrc.bio.indiana.edu//stock/1667227; RRID:DGRC_1667227)) was obtained from the Drosophila Genomics Resource Center. The cDNA was already in the pGW-HA.attB vector backbone for expression under the UASt regulatory sequence. We performed mutagenesis on the wildtype (WT) cDNA using Q5 PCR to generate two microcephaly patient mutations: c.182_187del, and c.1235_1236insA. The primers used to generate these mutations are listed in the key resources table. All cDNA vectors were injected by BestGene Inc. into VK37 flies that contain an attP docking site at 22A3 on 2L and transgenics were generated using phiC31 integrase-mediated recombination into the second chromosome.

### Immunohistochemistry

All immunohistochemistry was performed on late third-, second-, 24-hour first-, and early first-instar larval brains. Brains were dissected in phosphate-buffered saline (PBS) and transferred to microcentrifuge tubes for a 20-minute fixation with 4% paraformaldehyde in PBS + 0.3% Triton X-100 (PBST) and then washed with PBST three times for five minutes. Next, the brains were washed twice with PBST + 5% Bovine serum albumin (PBSTB) for 30 minutes, followed by a 30-minute block with PBSTB + 5% Normal Donkey Serum. The brains were then incubated in primary overnight at 4°C. The following primary antibodies were used and category numbers are listed in the key resources table: rat anti-Deadpan (neural stem cells, 1:1000), rabbit anti-pHH3 (proliferation, 1:500), rat anti-Miranda (neural stem cells, 1:1000), rabbit anti-GFP (1:1000), and mouse anti-Prospero (1:1000, early neurons).

The next day, brains were washed three times with PBSTB for 20 minutes and incubated in PBSTB + secondary antibodies for one hour. The following secondaries were used at 1:500 Donkey anti-rat Alexa fluor 647, Donkey anti-rabbit Alexa fluor 488, Donkey anti-rabbit Alexa fluor Rhodamine RedX, Donkey anti-mouse Rhodamine RedX, and 1:1000 DAPI. Finally, brains were washed 3 times with PBST and mounted in slow fade gold.

### Confocal microscopy

A singular brain lobe was imaged per brain. The following confocal settings were used to image: 40X water immersion lens, zoom of 0.7 for third-instar and 1 for first-and second-instars, frame size of 1024×1024, line averaging of 2, scan speed of 8, z-slice size of 2μm, and z limits were set using the Deadpan or Miranda channel to capture the entire brain lobe.

## QUANTIFICATION AND STATISTICAL ANALYSIS

For the following analyses, a singular brain lobe was analyzed per brain using the software system Imaris, and the statistics were performed using GraphPad Prism. Raw p-values are in the figures. A p-value of <0.05 was considered significant.

### Brain lobe volume

Using the surface function in Imaris, the perimeter of the brain was traced using the Deadpan channel at every 5^th^ z slice and then compiled to form a 3D surface. Volume was recorded from the statistics tab. An independent t-test was used for comparing brain lobe volume between control and *Bx42* knockdown for first and second instar brains. A multiple comparisons ANOVA was performed for both rescue experiments. In the *prospero* knockdown rescue, all conditions were compared to the *EGFP* RNAi; *Bx42* RNAi group. In the human rescue, all conditions were compared to the *UAS-Luciferase*; *Bx42* RNAi group.

### Proliferation

Using the spots function in Imaris, ventral central brain neural stem cells were manually counted. Next, the number of neural stem cells with pHH3 colocalized within the nucleus were counted. The percentage of dividing cells was then calculated by dividing the pHH3+ cells over total neural stem cells. An independent t-test was run for first-and second-instar brains. Tukey’s or Sidak’s multiple comparisons ANOVA was run for rescue experiments. In the *prospero* knockdown rescue, all conditions were compared to the *EGFP* RNAi; *Bx42* RNAi group. In the human rescue, all conditions were compared to the *UAS-Luciferase*; *Bx42* RNAi group.

### Percentage of nuclear Prospero

A similar procedure to determine the percentage of pHH3+ was used to determine the percentage of nuclear Prospero. Deadpan is a nuclear marker when the cell is not dividing, so if Prospero colocalized with Deadpan during interphase, it was considered nuclear Prospero. The number of ventral central brain neural stem cells with nuclear Prospero was counted manually. The number of nuclear Prospero+ cells was divided by the total number of ventral central brain neural stem cells to get the percentage of nuclear Prospero stem cells. An independent t-test was performed between control and *Bx42* RNAi for each larval stage.

### Protrusions

Protrusions were identified using the Miranda antibody. Miranda has been well-established for identifying protrusions by morphological characterization (34–36). Any neural stem cell that had a singular protrusion extending from the cell body towards the neuropil was counted by hand using the spots function in Imaris. On a separate spots tab, all neural stem cells were counted to determine percentage of all stem cells with protrusions. A multiple comparisons ANOVA was used to compare changes in experimental condition and developmental stage.

